# Effects of CXCL12/CXCR4/CXCR7 axis on human sperm motility and chemotaxis

**DOI:** 10.1101/831065

**Authors:** Chen Wang, Jiepin Huang, Ling Ding, Ruohan Huang, Luru Dai, Wenhui Zhou

## Abstract

Mammalian spermatozoa undergo a series of functional changes in the female genital tract before they can fertilize oocytes. Chemokines and their receptors are involved in these complex biological events. However, the detailed molecular mechanisms on the modulation of chemokines on spermatozoa still need to be elucidated. CXCL12, a chemokine, has been widely studied in leukocyte migration and. Here we found that CXCL12 was existent in human cumulus cells and follicular fluid and its receptors CXCR4 and CXCR7 were co-expressed on human spermatozoa. We investigated the effects of CXCL12 on various biological functions of human sperm and which receptor plays a dominant role in these processes. We found that CXCL12 could promote human sperm chemotaxis, motility, penetration in the mucus, acrosome reaction and Ca^2+^ influx through CXCR4 rather than CXCR7. In addition, the simplified physical model reasonably explained the change of sperm velocity under the influence of CXCL12 concentration gradient, which was identical to their physiological motion patterns.

## Introduction

Successful fertilization is the basis of embryo development and implantation. How do spermatozoa reach to and combine with the oocyte is a pivotal issue in the process of fertilization. It is generally recognized that when mammalian spermatozoa transit through the female genital tract to the cumulus-oocyte complex (COC), they must undergo a series of complex functional changes under the action of various maternal-derived substances before they can fertilize oocytes, collectively called capacitation (Sugiyama and Chandler, 2014; Puga Molina et al., 2018). The capacitated spermatozoa could be hyperactive and swim towards COC in oviduct with higher speed and asymmetrical oscillations under the propelling of the flagellum (Leemans et al., 2015; De Lisa et al., 2013). However, the mechanisms involved in sperm capacitation and why the spermatozoa can accurately arrive at the fertilization site, especially in human reproduction process, have not been clearly elucidated.

Chemotaxis is one of the most important factors in sperm guidance during fertilization among marine invertebrates, which provides a good inspiration to study the fertilization of human (De Lisa et al., 2013; Hussain et al., 2016). The sperm-attracting substances secreted by female genital tract are called chemoattractant. After binding with specific sperm surface receptors, chemoattractant activates downstream pathways, increase the intracellular cAMP concentration and Ca^2+^ influx, and accelerate sperm motility (Blengini et al., 2011; Lymbery et al., 2017). CXCL12 is a unique chemoattractant and plays an important role in the growth, metastasis and invasion of tumour cells (Esencay et al., 2013; Wang et al., 2015). Studies in sheep have shown that CXCL12 is involved in early placental angiogenesis, and loss of the CXCL12 gene is linked to the occurrence of preeclampsia and abortion (Quinn et al., 2014; Quinn et al., 2016). CXCL12 is also involved in the migration of early embryonic progenitor cells and gene regulation during follicular maturation (Sayasith and Sirois, 2014; Knaut et al., 2002; Molyneaux, 2003).

Our previous studies have verified the expression of CXCL12/CXCR4/CXCR7 at the maternal-fetal interface during menstral cycle and early pregnancy (Zhou et al., 2015; Zhou et al., 2008; Ren et al., 2012; Zheng et al., 2018) and found that CXCL12/CXCR4 signalling pathways not only in an autocrine way participate in the regulation of migration and invasion of human first-trimester trophoblast cells and endometrial epithelial cells, but also in a paracrine manner regulate the migration of decidual stromal cells and endometrial epithelial cells, which promotes the dialogue and exchanges between these cells and helps the formation of maternal-fetal interface of human early pregnancy (Zhou et al., 2008; Ren et al., 2012; Zheng et al., 2018). These results reveal an essential role of CXCL12/CXCR4/CXCR7 axis in human implantation and early pregnancy. However, the role of this axis in human fertilization process and its effect on human sperm biological function still remains an open question.

The aims of this work were first, to examine the expression of CXCR4, CXCR7 and CXCL12 in human spermatozoa and cumulus cells; and second, to determine the effects of CXCL12 on the modulation of human sperm function, including motility, chemotaxis, acrosome reaction and intracellular Ca^2+^ concentration. This study could improve the understanding of the mechanisms of changes of human sperm function in female genital tract before fertilization and provide new clues for the cause, diagnosis and treatment of unexplained infertility.

## Results

### Contents of CXCL12 in human cumulus cell and follicular fluid

Immunofluorescence staining shown in Fig. 1A revealed green staining specific for CXCL12 in the cytoplasm of human cumulus cells. As shown in the quantitative reverse transcription polymerase chain reaction (qPCR) results presented in Fig. 1B, CXCL12 and its receptor genes were transcribed in human cumulus cells, and the mean levels of CXCL12, CXCR7 and CXCR4 were 24.77, 84.97 and 879.09, respectively. As shown in Fig. 1C, the mean concentration of CXCL12 in human follicular fluid was 2545.34 pg/ml.

**Figure 1:**
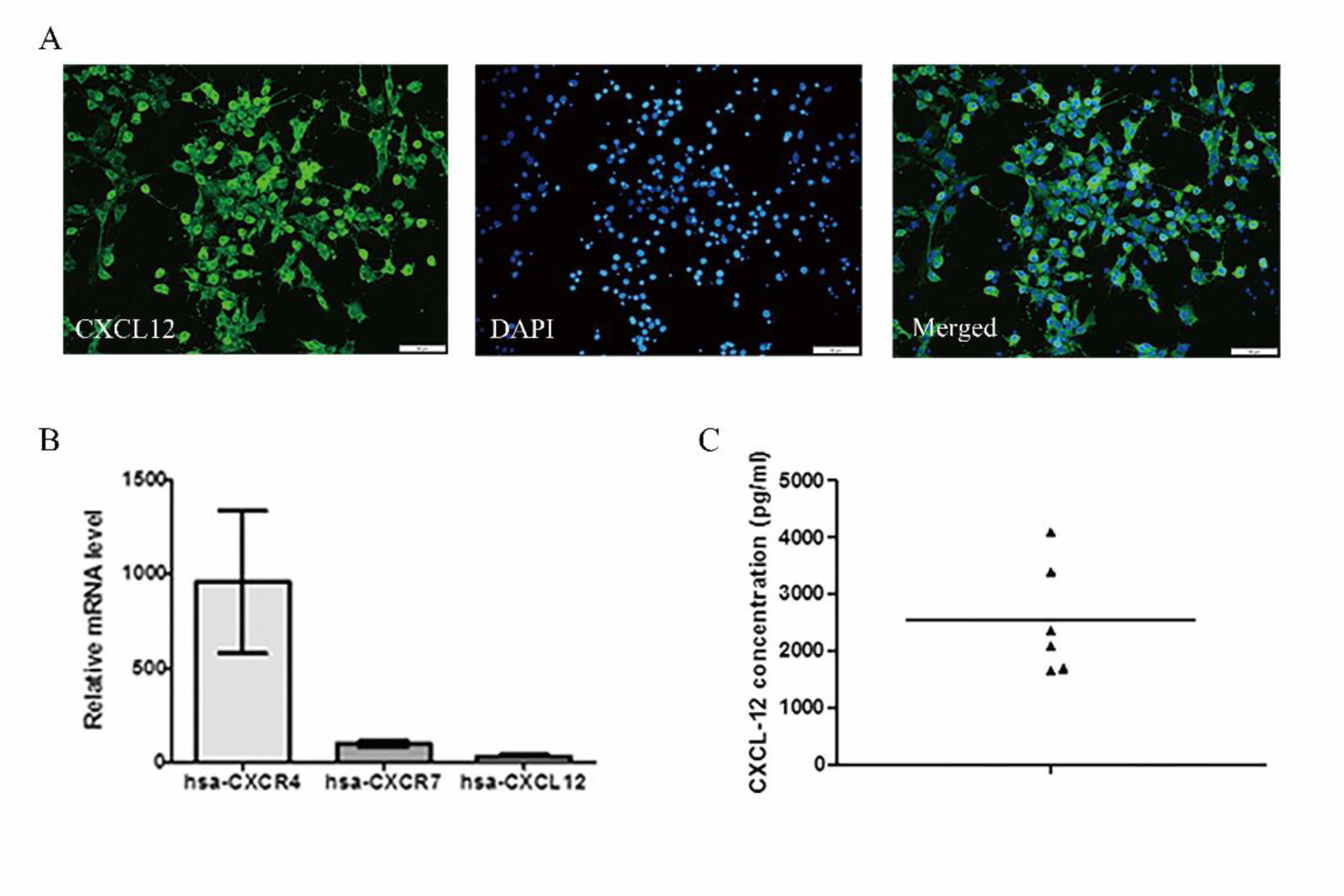
Identification and expression of CXCL12 in human cumulus cells. (A) Indirect immunofluorescence revealed an expression of CXCL12 (green) in human cumulus cells. Magnification: × 400. Experiments were performed using 5 independent samples. Bar was 50 μm. (B) CXCR4, CXCR7 and CXCL12 mRNA relative expression levels in cumulus cells. GAPDH was used as an internal control. Experiments were performed using 7 independent samples. (C) ELISA analysis was performed to quantify the amount of CXCL12 in 6 independent samples of human follicular fluid.

### Expression of CXCR4 and CXCR7 in human spermatozoa

Immunofluorescence showed that CXCR4, the receptor of CXCL12, was expressed in the midpiece and flagella of spermatozoa, while CXCR7 was widely distributed on the surface of the sperm membrane (Fig. 2A). The protein levels of CXCR4 and CXCR7 were detected by western blotting. CXCR4 and CXCR7 had positive expressions, but CXCL12 was barely detected. The relative intensities of the CXCR4 and CXCR7 were 1.683 ± 0.2251 and 1.7025 ± 0.0238 respectively.

**Figure 2:**
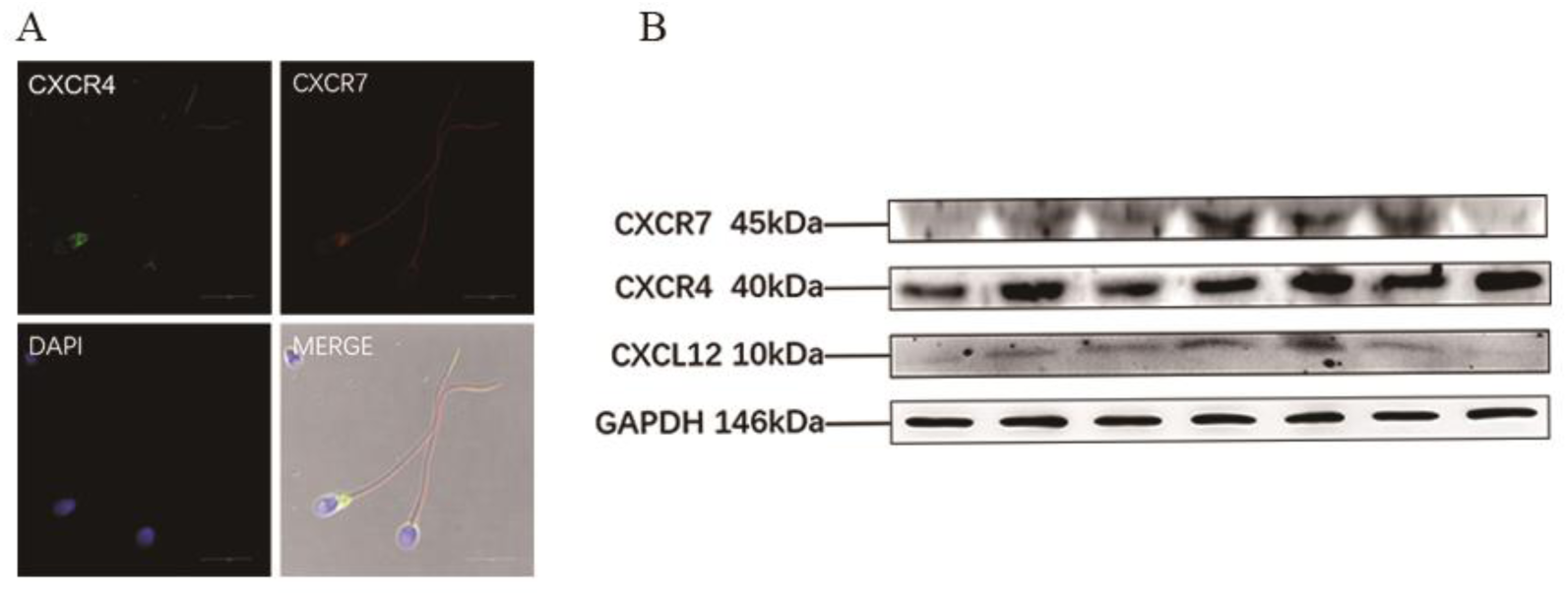
CXCR4 and CXCR7 were located in human spermatozoa. (A) Indirect immunofluorescence was used to detect the localization of CXCR4 and CXCR7 in human spermatozoa. CXCR4 (green) was mainly restricted in the post-head, midpiece and the rear of the flagellum. CXCR7 (red) was spread widely on the whole sperm membrane. Magnification: × 1000. Experiments were performed using 5 independent samples. The scale bars were 20 μm. (B) A representative western blot picture of CXCR4 and CXCR7 expression in human spermatozoa. Experiments were performed using 7 independent samples. CXCL12 protein was not detected and β-actin was used as a control.

### CXCL12 promoted chemotaxis of spermatozoa via CXCR4

The trans-well assay is a common method to study sperm chemotaxis ^18^. In our experiments, we found that CXCL12 at 100 ng/ml and 10 μg /ml, respectively, could significantly attract spermatozoa (in 10^6^) to move to the bottom well (2.73 ± 0.123 vs. 3.34 ± 0.117; *P* = 0.003; and 2.73 ± 0.123 vs. 3.28 ± 0.065; *P* = 0.01). When blocking with AMD3100 at 100 ng/ml, the increased chemotaxis of sperm induced by CXCL12 decreased significantly (3.34 ± 0.117 vs. 3.01 ± 0.09; *P* = 0.022), while the anti-CXCR7 neutralizing antibody had no effect (3.12 ± 0.094 vs. 3.12 ± 0.09; *P* = 0.14). These results suggest that CXCL12 promotes chemotaxis of human spermatozoa via CXCR4 rather than CXCR7.

### Effects of CXCL12 on sperm motility

We found that 100 ng/ml CXCL12 could significantly improve the curvilinear velocity (VCL) of sperm (71.00 ± 1.64 μm/s vs 77.17 ± 1.04 μm/s; *P* = 0.027) while other motility parameters were not affected after various concentrations of CXCL12 treatment (Fig. 4A). Therefore, we explored the roles of CXCR4 and CXCR7 in sperm velocity, and whether CXCR4 and CXCR7 could respond to CXCL12 stimulation simultaneously. AMD3100 is a specific inhibitor of CXCR4, which significantly decreased the improvement of sperm VCL produced by 100 ng/ml CXCL12 (77.17 ± 1.04 μm/s vs. 61.33 ± 1.89 μm/s; *P* = 0.000). The neutralizing antibody of CXCR7 failed to affect the motility parameters of spermatozoa with or without CXCL12 addition. Interestingly, we found that AMD3100 decreased the motility of spermatozoa even without stimulation by CXCL12 (VCL 71.00 ± 1.64 μm/s vs. 58.32 ± 1.73 μm/s, *P* = 0.025; VSL (strait-line velocity) 43.45 ± 1.16 μm/s vs. 30.43 ± 0.75 μm/s, *P* = 0.000; VAP (average path velocity) 47.39 ± 1.11 μm/s vs. 33.85 ± 0.91 μm/s, *P* = 0.00; ALH (amplitude of lateral head displacement) 5.57 ± 0.36 μm vs. 4.41 ± 0.23 μm, *P* = 0.002; Fig, 4B).

**Figure 3.**
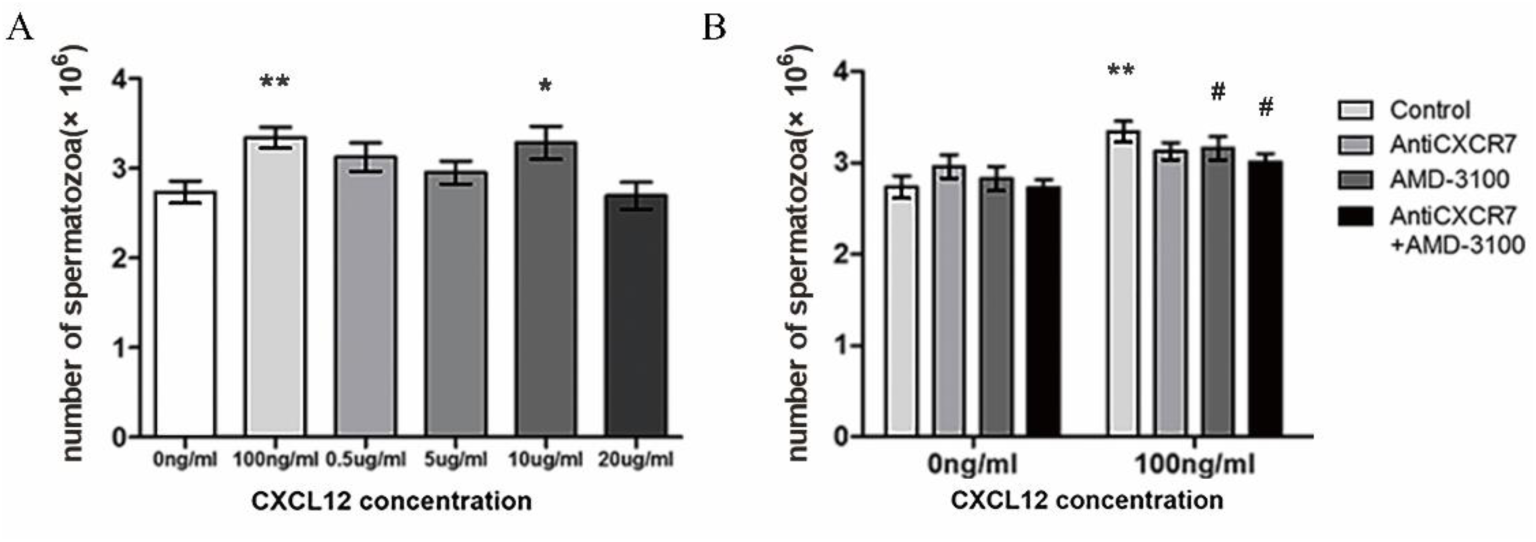
Concentration-dependent profile of sperm chemotaxis elicited by CXCL12/CXCR4. (A) This graph illustrated the numbers of spermatozoa (in 10^6^) swimming towards different concentrations of CXCL12 (n = 8). (B): Samples pre-treated with AMD3100 (10 μΜ), CXCR7 antibody (20 μg/ml) or recombinant human CXCL12 (100ng/ml). CXCR7 antibody alone had no effect on sperm chemotaxis while AMD3100 obviously decreased the chemotaxis induced by CXCL12 (n = 13). Bars indicated the mean ± SEM. Compared to the corresponding control, \**P* < 0.05 and ** *P* < 0.01. Compared to the corresponding CXCL12 treatment group, #*P* < 0.05, ##*P* < 0.01.

**Figure 4:**
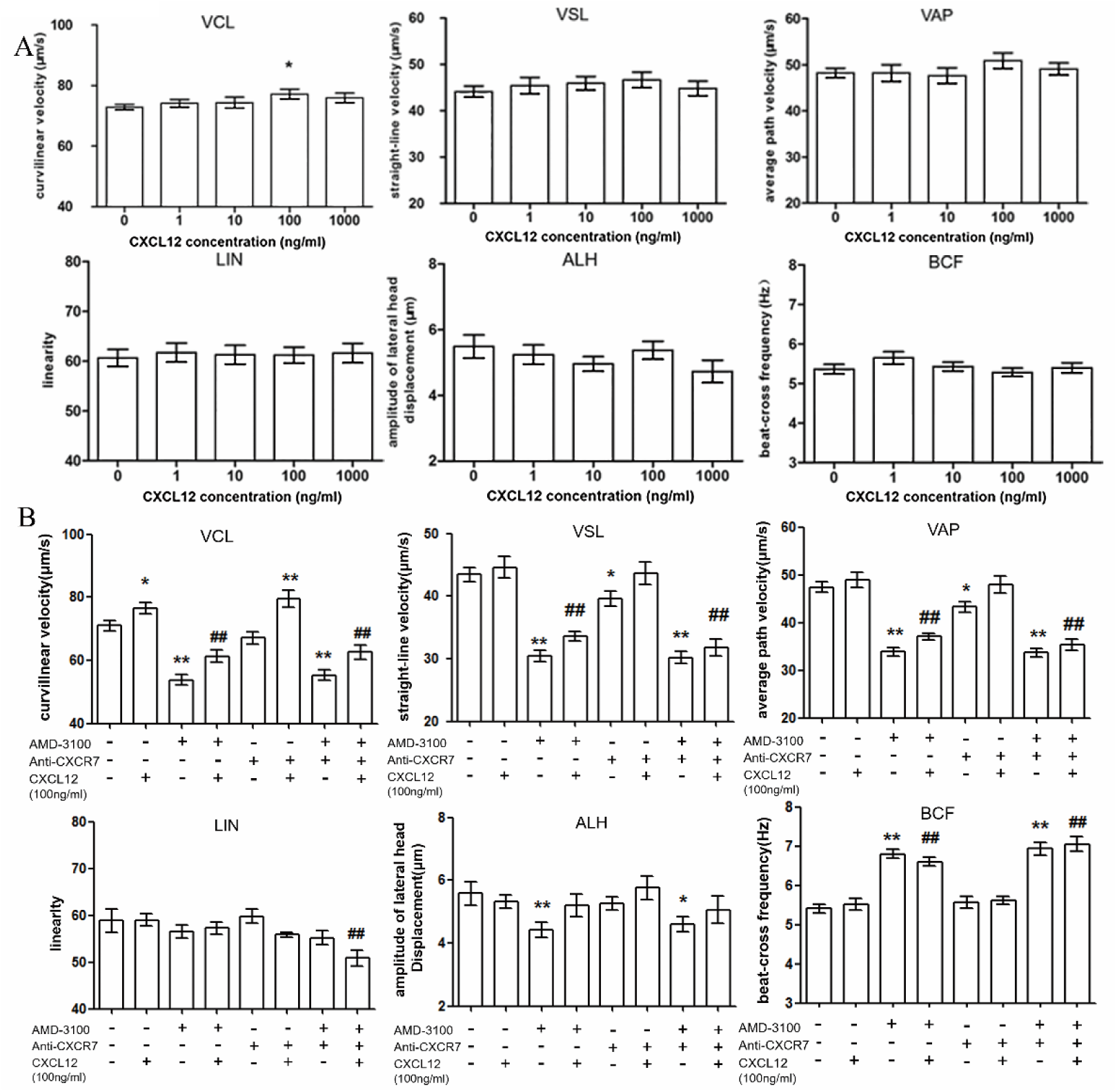
Effects of CXCL12/CXCR4/CXCR7 on human sperm motility parameters. (A) After stimulation with different concentrations of CXCL12, a significant increase of sperm VCL at 100 ng/m of CXCL12 was observed with CASA analysis (n = 8). Various concentrations of CXCL12 had no effect on other motile parameters of sperm. (B): Sperm samples were treated with CXCR4 inhibitor AMD3100 (10 μΜ), anti-CXCR7 antibody (20 μg/ml), recombinant human CXCL12 (100 ng/ml) or a combination of them (n = 16). AMD3100 significantly decreased the velocity of spermatozoa with or without CXCL12 addition (n = 8). Bars indicated the mean ± SEM. Compare to the corresponding control, \**P* < 0.05 and \*\**P* < 0.01. Compared to the corresponding CXCL12 treatment group, #*P* < 0.05 and ##*P* < 0.01 CASA: computer assisted sperm analysis, VCL: curvilinear velocity, VSL: strait-line velocity, VAP: average path velocity, LIN: linearity ALH: amplitude of lateral head displacement, BCF: beat across frequency.

### Simplified physical model for sperm motility

In the liquid environment, the sperm motility is quite different from the motion of macroscopic objects in our daily life. When sperm moves slowly in viscous fluid such as seminal plasma, the inertia force of the fluid is negligible in comparison with the viscous frictional force. We call this liquid environment a low-Reynold-number environment.

A large number of experiments show that the trajectory of free-swimming sperm is curved. That is to say, sperm trajectory will be a circle without considering the influence of random forces of itself and environment. This curvilinear motion originates from the asymmetric swing of flagellum, which deflects the direction of sperm movement.

In the resting state, sperm makes a circular motion due to the asymmetric swing of flagellum. When sperm senses chemoattractant concentration, a series of cascade reactions are triggered, which leads to the increase of intracellular concentration of Ca^2+^, and aggravates the asymmetry of flagella oscillation. As a result, the curvature of sperm trajectory increases and the chemotaxis turns. This phenomenon has also been found in a large number of experiments ^19^. Now we hope to propose a physical model to explain the effect of chemoattractant concentration on sperm motility velocity. We simplify the head of sperm into a rigid sphere (as shown in Figure 5). The dotted line in the figure is the central axis of the ellipsoid head of sperm. We assume that the velocity of sperm is in this direction every moment. *F* is the force generated by the flagellum. *θ* is the angle of the force departed from the central axis.*r* represents the size of the sperm.

**Figure 5.**
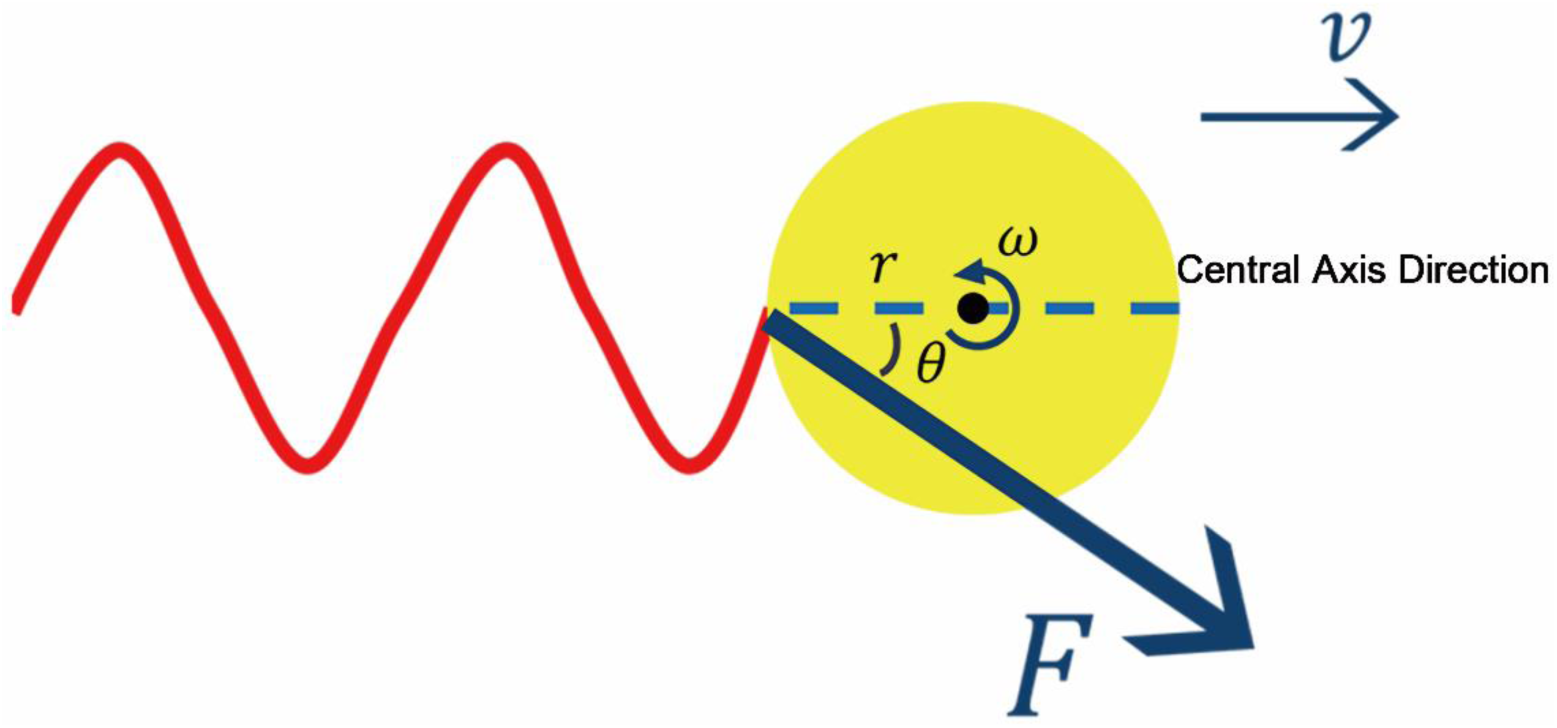
A simplified physical model of sperm motion. We simplified the head of sperm into a rigid sphere. The dotted line in the picture was the central axis of the ellipsoid head of sperm. We assumed that the speed of sperm was in central axis direction every moment.

The physical reason for sperm motility is that every point on the flagellum exerts a force on the fluid, which will give the flagellum a reaction force. Under the action of this reaction force, the swing of flagellum causes the forward movement of sperm. If we divide the sperm into two parts, the head and the tail flagellum, then the force of the flagellum on the head will act at the junction of the two parts, as shown in the figure above. It should be noted that the force exerted on the head by the flagellum does not necessarily follow the direction of the central axis, because the average axis of the flagellum after removing the sinusoidal waveform does not necessarily follow the direction of the central axis, which is caused by the asymmetry of flagellum swing. It is precisely because the force does not follow the direction of the central axis that makes the head rotate, thus triggering the circular movement of sperm.

We can resolve the force of flagellum to the head into two directions: along the axis direction and perpendicular to the axis direction. Using the formulas of forces and torques in the low-Reynold-number environment, we can obtain

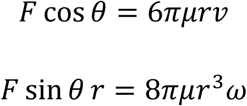

where *μ* represents the viscosity coefficient of fluid environment. These equations mean that the force along the central axis causes the sperm to move forward at the speed *v*, and the force perpendicular to the central axis generates the torque, which provides the angular velocity *ω*, so that the head can rotate, thus resulting in a circular motion.

From the above two equations we obtain the speed of sperm

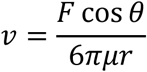

and the trajectory curvature

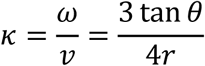

On the one hand, we see that curvature *κ* depends on angle *θ*. According to the phenomena observed in our experiments, the greater the concentration, the greater the curvature *κ*, which also requires the larger the angle *θ*. Then cos *θ* will decrease, *v* should decrease if *F* is unchanged.

On the other hand, Figure 4 (beat across frequency, BCF) shows that the higher the concentration, the greater the frequency of flagellar oscillation. This is because higher ligand concentration leads to higher ligand-receptor binding probability and the cascade reactions are triggered more strongly. So *F* should also increase; as a result, if *θ* is constant, *v* should increase.

In the above discussions, we find that there are two consequences when the chemoattractant concentration increases: one is that *F* becomes larger, the other is that *θ* becomes larger. We can see, in the expression of sperm motility velocity, they have opposite effects on the velocity *v*. This leads to the phenomenon that the speed of sperm increases first and then decreases with the increase of chemoattractant concentration.

### Sperm penetration in artificial viscous medium

Flattened capillary tubes filled with 1% methylcellulose were used *in vitro* to simulate the viscosity of female genital fluids ^20^. CXCL12 of 100 ng/ml could significantly improve the penetration ability of sperm (14.93 ± 0.605 vs. 17.21 ± 0.81; *P* = 0.042; Fig. 6A), which was significantly inhibited by AMD3100 addition (17.21 ± 0.813 vs. 13.25 ± 0.97; *P* = 0.002). The penetration of sperm in the AMD3100 treatment alone group was significantly reduced either compared to corresponding group (14.93 ± 0.605 vs. 11.06 ± 0.63, *P* = 0.000). The CXCR7 neutralizing antibody had no effect on the penetration of human spermatozoa in methylcellulose (Fig. 6B).

**Figure 6.**
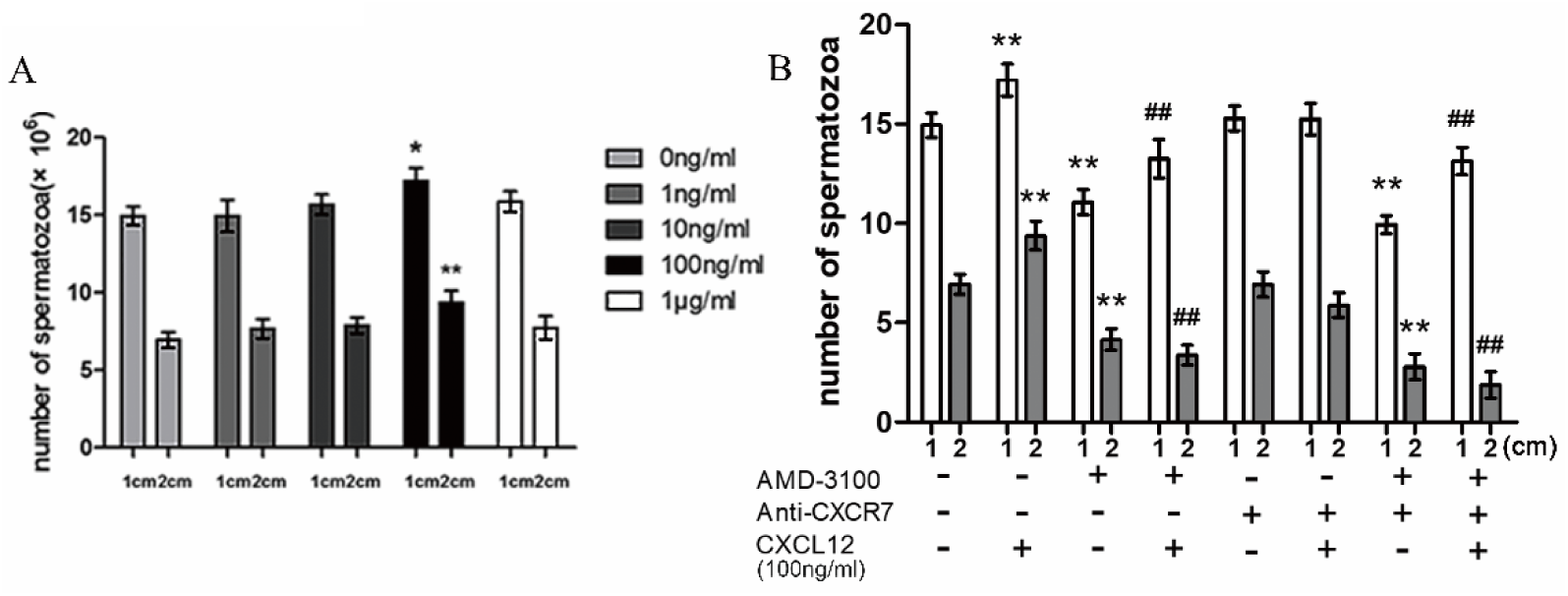
CXCL12/CXCR4/CXCR7 affected the penetration of spermatozoa through artificial viscous medium. (A) Capacitated sperm samples were stimulated with different concentrations of CXCL12 (n = 8). CXCL12 of 100 ng/ml obviously improved the penetrating ability of spermatozoa. (B) Capacitated sperm samples were treated with different combinations of AMD3100 (10 μΜ), anti-CXCR7 antibody (20 μg/ml) and recombinant human CXCL12 (100 ng/ml). A significant decrease in the sperm numbers at both 1 and 2 cm was observed after addition of AMD3100 with or without CXCL12 pre-stimulation (n = 16). Bars indicated the mean ± SEM. Compared to the corresponding control, \**P* < 0.05, \*\**P* < 0.01. Compared to the corresponding CXCL12 treatment group, #*P* < 0.05, and ##*P* < 0.01.

### CXCL12 increased the level of intracellular Ca^2+^ and the incidence of acrosome reactions in human spermatozoa

Previous studies have found that chemokines affect sperm function by changing intracellular Ca^2+^ concentrations (Guerrero et al., 2010; Jaldety and Breitbart, 2015). Here we found that 100ng/ml CXCL12 stimulation significantly increased the intracellular Ca^2+^ concentrations in spermatozoa (1.17 ± 0.022 vs. 1 ± 0.05; *P* = 0.001; Fig. 7A). Treatment with AMD3100 significantly reduced the intracellular concentration of Ca^2+^ (0.956 ± 0.031 vs. 1 ± 0.05; *P* = 0.047) induced by CXCL12 addition. However, after blocking CXCR7, the Ca^2+^ level was not affected (1.011 ± 0.029 vs. 1 ± 0.059; *P* = 0.836; Fig. 7B). Ca^2+^ level significantly decreased after treated with AMD-3100 even without CXCL12 stimulation (0.885 ± 0.023 vs. 1 ± 0.059; P = 0.836). NNC 55-0396 is a highly selective T-type calcium blocker. In this study, it was used to investigate changes in Ca^2+^ in sperm influx from human tubal fluid (HTF; ART-1020) medium. We found that exogenous CXCL12 did not change intracellular calcium concentration after NNC pre-treatment (0.945 ± 0.021 vs. 0.894 ± 0.018; *P* = 0.107).

**Figure 7.**
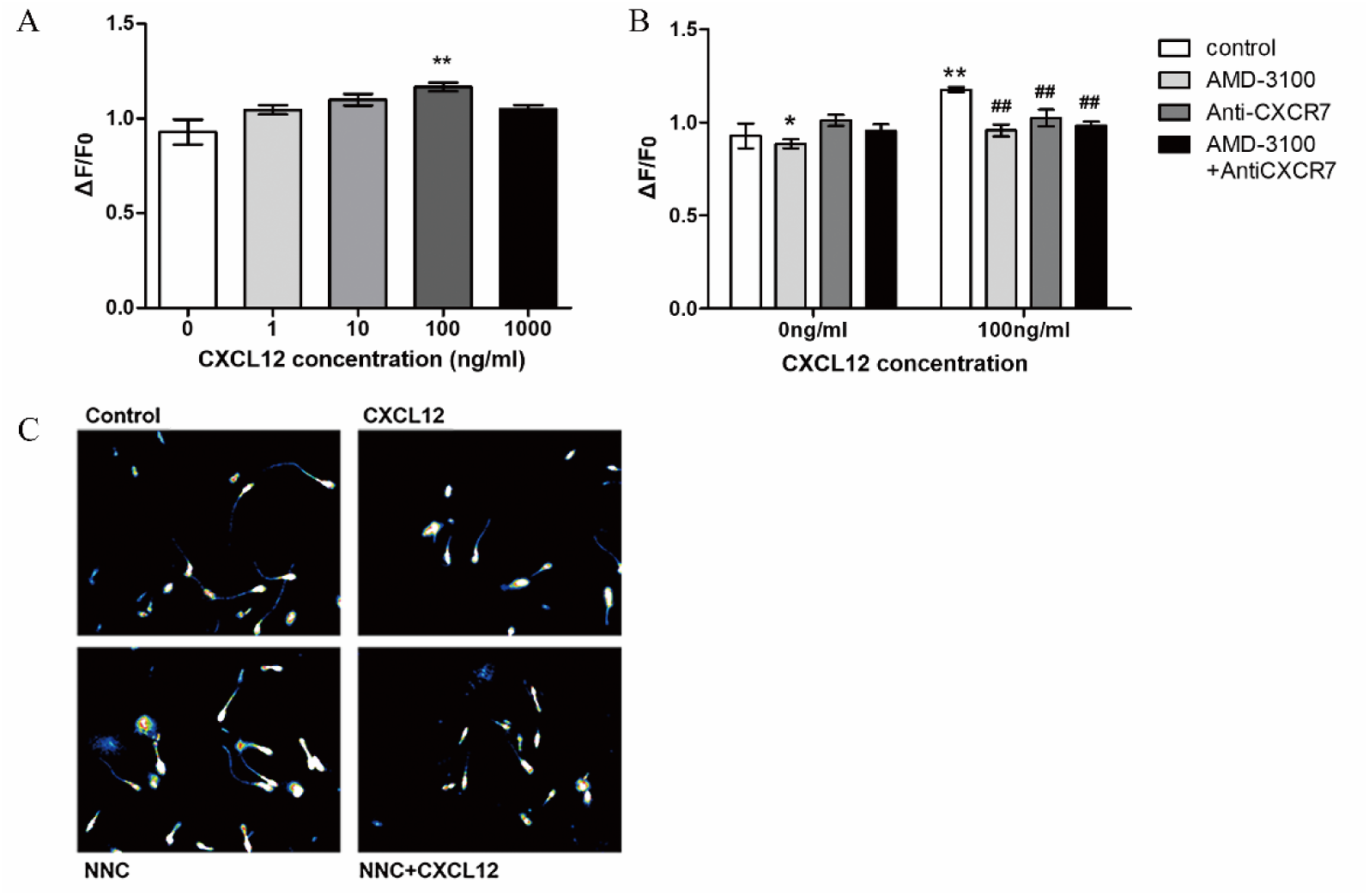
Effects of CXCL12/CXCR4/CXCR7 on sperm intracellular Ca^2+^ concentration. (A) Comparison of mean calcium fluorescence intensity ratio (ΔF/F_0_) in the sperm head and midpiece after treating with different concentrations of CXCL12 (n = 10). F_0_ represented the fluorescence baseline of each sample with no stimulus; F represented the average fluorescence intensity of the various treatments. (B) Sperm calcium fluorescence in AMD3100 (10 μΜ) and CXCR7 antibody (20 μg/ml)-treated samples in response to CXCL12 at 100 ng/ml (n = 8). (C) Representative micrographs of spermatozoa with or without 1 μM NNC 55-0396 incubation (left panel) or supplemented with 100 ng/ml CXCL12 for 5 min in HTF medium (n = 4). Bars indicated the mean ± SEM. Compared to the corresponding control, \**P* < 0.05, ** *P* < 0.01. Compared to the corresponding CXCL12 treatment group, #*P* < <0.05, ##*P* < 0.01. HTF: human tubal fluid.

In this study, Ca^2+^ ionophore A-23187 was used as a positive control for measuring the percentages of acrosome reactions ^23^. We found that CXCL12 treatment significantly increased the percentage of acrosome reactions compared with the untreated control samples (9.78 ± 0.73% vs. 5.67 ± 0.36%; *P* = 0.000; Fig .8A), and this effect was significantly reduced when jointly blocking CXCR4 and CXCR7 (4.60 ± 0.509%; *P* = 0.000) and when only blocking CXCR4 (5.24 ± 0.602%; *P* = 0.000; Fig. 8A).

**Figure 8.**
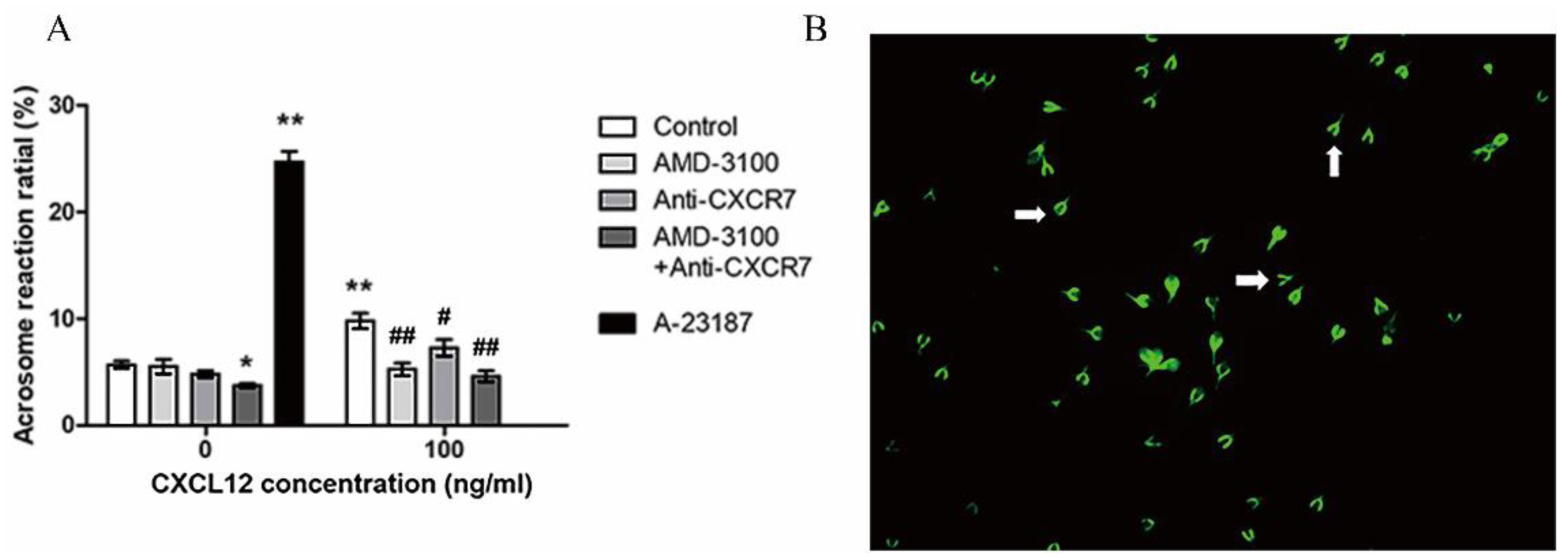
Effects of CXCL12/CXCR4/CXCR7 on acrosome reaction. (A): Acrosome reactions (%) induced by CXCL12. The acrosome reaction rate significantly decreased after blocked with AMD3100 (10 μΜ) or CXCR7 neutralizing antibody (20 μg/ml). A-23187 (10 μΜ) was used as a positive control (n = 6). (B) A typical figure of acrosome reactions induced by A-23187. Arrows showed the sperm with acrosome reactions. Compared to the corresponding control, \**P* < 0.05, ** *P* < 0.01. Compared to the corresponding CXCL12 treatment group, #*P* < <0.05, ##*P* < 0.01.

## Discussion

The complex mechanism why only a few outdoing hundreds of millions of spermatozoa in every ejaculation finally get access to COC in human fertilization still remains uncertain ^24^. As a necessary guidance condition for aquatic sperm to find oocyte successfully, chemotaxis has attracted wide attention^25^. Currently, stable and persistent chemokine guidance in female genital tract is considered as an important prerequisite for sperm to locate in the COC correctly^26^. Follicular fluid and cumulus granulosa cells can secrete a variety of chemokines, which promote sperm hyperactivity and improve sperm motility following receptor binding (Tacconis et al., 2001;Villanueva-diaz and D, 1990). However, ethical and experimental factors as well as the inability to observe the fertilization process directly have limited our insight into the contribution degree of various chemokines and their interactions during human fertilization process.

Though extensively reviewed in leukocyte migration (Guyon, 2014;Vitiello et al., 2016), the role of CXCL12 in human fertilization has not been verified. Some studies have found that there was a certain concentration of CXCL12 in female genital tract lavage fluid ^30^ and our previous study have demonstrated the persistent secretion of CXCL12 by endometrial epithelial cells ^17^. Furthermore, in the present study, we also confirmed that CXCL12 was expressed in human follicular fluid and cumulus granulosa cells, and its receptors CXCR4 and CXCR7 were expressed on sperm surface. Therefore, this study speculated that the existence of CXCL12 in female genital tract might affect sperm function after binding with its receptor.

We first detected the chemotaxis of CXCL12 on sperm with trans-well method. It was found that the number of down-swimming sperms significantly increased after CXCL12 stimulation and blocking CXCR4 but not CXCR7 specifically eliminated the chemotactic attraction of CXCL12 to sperm, suggesting a role of CXCR4 and its downstream pathways in CXCL12 chemotaxis during human fertilization.

Furthermore, the results of sperm motility showed that only the VCL in all parameters increased significantly after CXCL12 treatment, which was consistent with the findings in other studies that not all kinetics parameters had significant changes (Sumigama et al., 2015;Armon and Eisenbach, 2011). As an important symbol of sperm motility, VCL is also a representative of hyperactivation (Rovasio, 1998; Suarez and Dai, 1992). The larger the VCL is within the unit time, the longer the sperm’s motility track is, indicating that the sperm’s motility is stronger. Several results found that in physiological conditions, if ovulation has not occurred, sperm will adhere to the storage area of the ampullary epithelium of the fallopian tube, and will resume kinetic energy after ovulation, which needs a high speed of spermatozoa movement to help them get out of the storage area and continue to move forward (Eisenbach et al., 2015;Demott and Suarez, 1992). Basing on our experimental results, we established a physical model to describe the sperm motility in female genital tract and to explain the phenomenon found by Eisenbach and Demott (Fig. 5), which is similar to the physiological movement pattern suggested by Alvarez ^19^. This physical model helps us qualitatively understand how the change in concentration affects the velocity. That is, with the increase of CXCL12 concentration, the sperm motility rate increased in a concentration-dependent manner and gradually decreased after reaching a higher concentration (100 ng/ml), while the BCF increased continuously. This movement mode is consistent with the sperm movement mode under physiological conditions. After entering the reproductive tract, it is necessary to continuously accelerate to overcome the resistance of mucus and promote the sperm to move forward (Suarez and Dai, 1992;Demott and Suarez, 1992). After reaching the vicinity of the COC, the sperm’s movement mode changes from forward movement to “drilling-top” movement with continuous head swing, in preparation for penetrating the zona pellucidum.

Moreover, we also investigated the effects of CXCL12 on sperm acrosome reaction. In our experiment, 100 ng/ml CXCL12 significantly increased the acrosome reaction rate of sperm without zona pellucida induction in vitro. There was no need for continuous calcium ion influx in the process of acrosome reaction, since a small amount of calcium ion influx triggered continuous release of calcium ion into the acrosome--an important calcium depot in sperm cells (Lievano et al., 1985;Costello et al., 2009;Herrick et al., 2005). Thus, we suggested that when COC stayed in the ampulla of fallopian tube, locally-formed high concentration of CXCL12 contributed to the acrosome reaction rate, and the last “obstacle” of transparent zone can be penetrated by sperm to finish the process of fertilization.

The effect of chemokines on sperm motility and acrosome reaction depended on the increase of intracellular calcium concentration (Brokaw et al., 1974; Wiesner et al., 1998). Calcium ions changes the beating pattern of the sperm flagellum by activating adenosine cyclase, which increased the synthesis of cAMP ^41^. It is accepted that calcium ions in sperm cells mainly came from two parts: the release of the stored calcium ions and the influx of calcium ions from extracellular fluid (Putney, 1986; Bedu-Addo et al., 2007). Catsper is an important specific calcium channel mainly located on the midpiece of sperm to mediate calcium influx(Alasmari et al., 2013; Lishko et al., 2012; Chung et al., 2014). We found that CXCL12 significantly increased sperm calcium concentration. In order to further identify the source of calcium ions, we used Catsper specific blockers and the results showed that CXCL12 had no significant effect on the change of intracellular calcium concentration in spermatozoa, reflecting that CXCL12 might regulate sperm motility by mediating extracellular calcium influx. However, the effect of calcium ion level changes on sperm motility and whether CXCL12 affects the release of intracellular calcium storage pools need further experimental investigations.

In our experiments, we found that AMD-3100, a specific blocker of CXCR4, could significantly reduce the kinetic energy and intracellular calcium concentration of sperm. CXCR4 was mainly expressed in the neck of sperm, where the specific calcium channel Catsper and the mitochondrial sheath were located concurrently, suggesting that CXCR4 may be related to the function of Catsper (Sugiyama and Chandler, 2014; Miller and Mayo, 2017), but the molecular mechanism needs to be further explored. Except for Catsper, there are some other plasma membrane calcium pumps on sperm cell membrane (Schuh et al., 2004; Okunade et al., 2004). Thus, whether there is an interaction between CXCR4 and these calcium pumps is still an open question.

In the other hand, specific blockade of CXCR7 had no significant effect on sperm motility or calcium concentration. Current studies suggested that although both CXCR4 and CXCR7 were CXCL12 receptors, they might have different biological characteristics in different cells, and be regulated by complex interactions (Krikun, 2018; Hattermann and Mentlein, 2013). In some cases, CXCR7 alone was enough to mediate the metastasis and infiltration of tumor cells after binding with CXCL12 through the activation of the downstream pathways (Sánchez-Martín et al., 2013; Qian et al., 2018). While in Madin-Darby canine kidney (MDCK) cells, CXCR7 played as a special scavenger of CXCL12 due to its similar structure to G protein-coupled receptor (Pluchino et al., 2018; Naumann et al., 2010). In our study, contrary to CXCR4, CXCL12 remarkably increased the VCL of sperm after specifically blocking CXCR7, thus we speculated that the function of CXCR7 in human sperm might be similar to that in MCDK cells and that it affected CXCL12 binding to CXCR4 by regulating the concentration of CXCL12. Of course, this hypothesis needs further studies.

According to experimental results, we proposed a model of the mechanism of CXCL12/CXCR4/CXCR7 in human fertilization process shown in Fig 9: some sperm pass through cervix and enter uterine cavity after ejaculation, and the capacitated sperm continues to move under the continuous action of low concentration CXCL12 in uterine cavity via CXCR4 ligation, as well as CXCR4 also mediates the calcium ion influx, which can further improve the movement speed and penetration ability of sperm. After ovulation, a small amount of follicular fluid and cumulus granulosa complexes could enter the fallopian tube. The CXCL12 and other chemokines secreted by COC would form a high concentration area in the ampulla, which attract sperm to swim to COC and trigger acrosome reaction after its exposure to COC, finally completing sperm-ovum binding.

**Figure 9.**
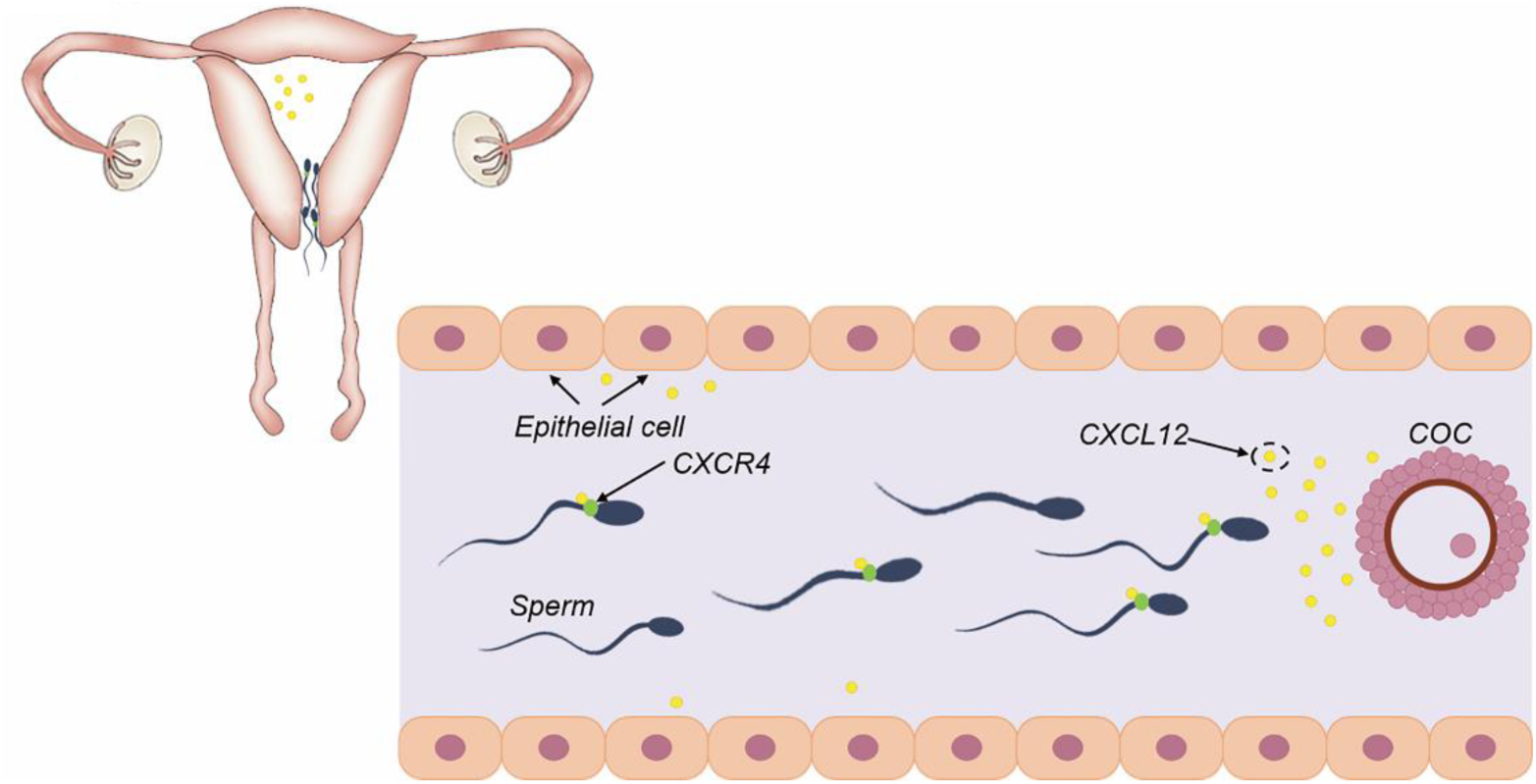
A model of CXCL12/CXCR4 involved in human fertilization process in female genital tract. A constant concentration of CXCL12 in the female reproductive tract guided sperm to swim toward the location of COC, during which CXCR4 highly expressed sperm made more responses.

There are still some questions needing further experimental exploration. Firstly, we still do not know whether CXCL12 could reduce the aimlessly curvilinear motion of sperm and make a more directional swimming, which can be tracked by 3D real-time trajectory. Secondly, the relationship between CXCR4 and the relative location as well as the downstream pathway of Catsper also need further investigation.

This study suggests that CXCL12 in female reproductive tract has a sustained and stable chemotaxis effect on sperm. CXCL12 mainly binds to the receptor CXCR4 instead of CXCR7 to promote calcium influx and regulate sperm movement. The physical model we established further elaborated the sperm movement pattern under the action of CXCL12, which could better understand the complex fertilization process. At the same time, our study may have potential clinical significance for the treatment of infertility and contraception.

## Methods

### Semen sample collection and preparation

Semen samples were collected from male partners among couples treated with in-vitro fertilization (IVF) for infertility caused by female factors, by masturbation after abstinence for 5 days. Samples were allowed to liquefy at room temperature for 30 min in sterile containers, then liquefied semen samples were first centrifuged at 1,500 *g* for 20 min, after which the sediments were recentrifuged at 10,000 *g* for 10 min to remove cellular elements and debris. Recovered spermatozoa were capacitated for 1 h in HTF (ART-1020) medium under 6% CO_2_ in humidified air at 37 °C. Semen parameters were consistently superior to WHO 2010 reference values for normal semen, and leukocytes were not found in any of the analysed samples.

Follicular fluid samples were collected from women who had undergone oocyte retrieval by transvaginal puncture for male factor infertility. The collected follicular fluid was centrifuged at 1500 *g* for 10 min. The supernatants were recentrifuged for 1000 *g* for 10 min, then stored at −80 °C.

Cumulus cells were derived from patients treated with intracytoplasmic sperm injection (ICSI) for male factor infertility. They were removed from COCs by mechanical and enzymatic methods before ICSI. After washing twice in HTF medium by centrifugation (1000 *g* for 5 min) cells were resuspended in Dulbecco’s Modified Eagle’s Medium (DMEM) containing 10% foetal bovine serum (FBS) and cultured at 37 °C under 6% CO_2_ in humidified air.

### Immunofluorescence

The concentration of spermatozoa was adjusted to 2 × 10^6^/ml with HTF medium, and air dried, while cumulus cells were cultured in four-well plates for 12 h. Samples were fixed in 4% formaldehyde at room temperature for 30 min, permeabilised with 0.02% Tween-20 for 30 min and then washed with phosphate-buffered saline (PBS) three times and blocked with PBS containing 600 μg/ml bovine serum albumin (BSA) for 1 h at room temperature. The slides were incubated overnight at 4 °C in a humidified chamber with primary antibodies: rabbit anti-human CXCR7 (Abcom ab72100), mouse-anti-human CXCR4 (MAB 172), or mouse-anti-human CXCL-12 (Abcom ab9797) diluted in blocking buffer (1:100). After washing in phosphate buffer saline with 0.1% Tween-20 (PBST) three times, the slides were incubated with secondary antibodies: donkey anti-rabbit IgG (ab 6800) or goat anti-mouse IgG (ab 6785) at 37 °C for 1 h. Finally, samples were incubated with 4’,6-diamidino-2-phenylindole (DAPI) (Solar-bio (S2110)) for 10 min at room temperature. Images were captured with an Olympus fluorescence microscope (Olympus, Tokyo, Japan). The experiment was performed with 5 independent samples.

### Western blotting

Precooled protein extracts were prepared after mixing protease inhibitors and 1 mM RIPA buffer with the samples. After 20 min incubation reaction on ice, they were centrifuged at 13,000g for 20 min at 4 °C, and the supernatant was retained. The protein concentration was adjusted to approximately 1 mg/ml with RIPA buffer and boiled for 5 min for denaturation.

The protein samples were detected by sodium dodecyl sulfate polyacrylamide gel electrophoresis (SDS-PAGE) (separation gel 12% w/v, 160 V, for 90 min and concentration gel 5% w/v at 90 V for 20 min). The PAGE membrane was immersed in 3% BSA-Tris-Buffered Saline Tween-20 (TBST) and incubated for 30 min at room temperature. The primary antibody was diluted with 3% BSA-TBST: mouse-anti-human CXCR7 antibody (ab72100; Abcam; 1:500), rabbit-anti-human CXCR4 antibody (ab2090; 1:500), rabbit-anti-human CXCL12 (ab155090; 1:200). Samples were incubated for 10 min at room temperature and kept overnight at 4 °C, then washed with TBST five times, 3 min each time. The secondary antibody was diluted with 5% skimmed milk powder, goat anti-rabbit IgG (H + L) (1:10000) and incubated for 40 min at room temperature. Samples were then washed with TBST five times, 3 min each time. MultiScan-3 (Thermo) was used to expose and develop the film. β-actin was used as an internal reference, and the concentration of the primary antibody mouse-anti-β-actin antibody (s001) was 1:1000. The secondary antibody Goat-anti-mouse IgG (1:5000) was diluted with 5% skim milk powder-TBST and incubated for 2 h at room temperature. The experiment was performed by Beijing center for physical and chemical analysis company with 7 independent samples.

### Quantitative reverse transcription polymerase chain reaction (qPCR)

The qPCR procedure was according to our previous research ^17^. Briefly, after centrifugation, 1 ml TRIZOL (Thermo) was added and samples were frozen at −80 °C. RNA was extracted as described. The RT reaction was performed follow the instruction of Fast Quant RT kits (with gDNAase) (TIANGEN). The cDNA was amplified using a fluorescence ratio PCR instrument (Roche 96). The final volume of the solution was 20 μl, including the cDNA (1 μl), SYBR FAST qPCR Master Mix (2 ×; 10 μl), and the primers to be tested (0.6 μl). After 5 min of pre-cycling at 95 ℃, we used 40 cycles at 95 °C for 15 s, 40 cycles at 60 °C for 20 s and 40 cycles at 72 °C for 15 s. The experiment was performed by Beijing Yuan Quan Yi Ke Bio-Tech Co. with 7 samples. The primer sequences were as follows (Table I):

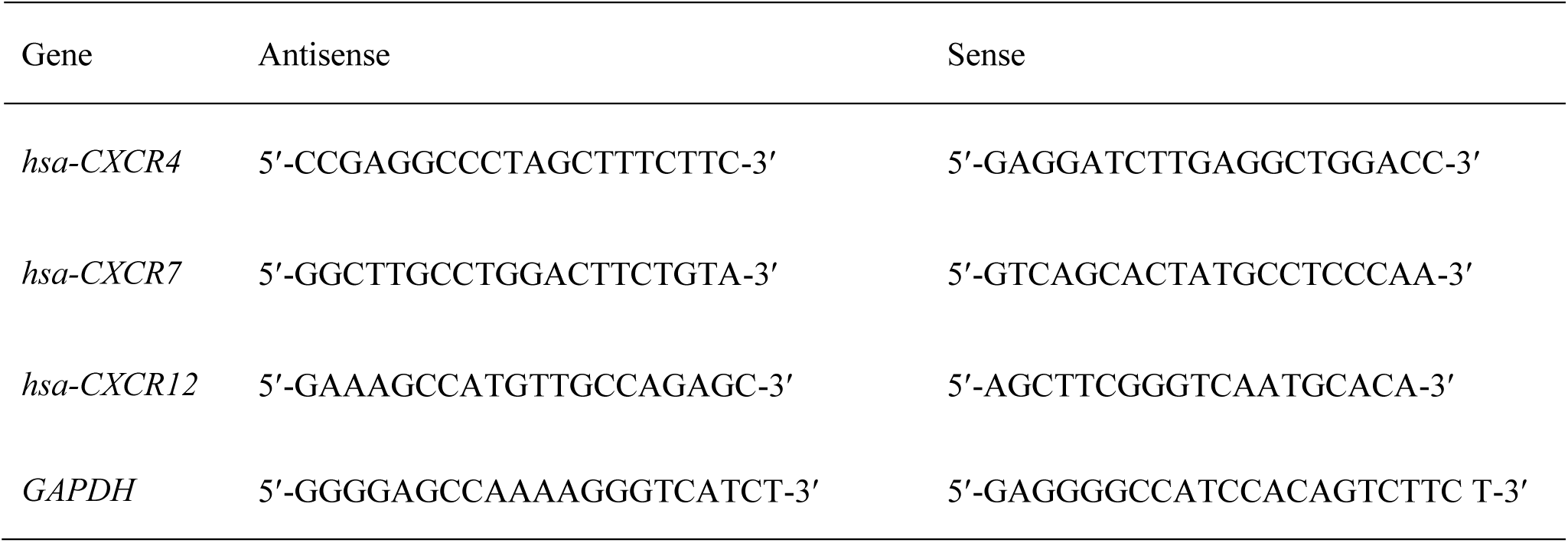

### Enzyme-linked immunosorbent assay (ELISA) protocols

Human follicular fluid samples were collected as described above. We used ELISA kits (Human CXCL12/SDF-1 alpha Quantikine ELISA, DSA00, R&D Systems) to explore whether cumulus granulosa cells secrete CXCL12, according to the manufacturer’s instructions and our previous publication ^17^. Briefly, the 96 well plates were placed on a microplate reader and light absorbance was read at 450-nm wavelength. The average value of each sample was used for statistical analysis. The ELISA was conducted in duplicate with 3 separate sample.

### Computer-assisted semen analysis

After capacitation, the sperm concentration was adjusted to 25-30 × 10^6^/ml with human tubal fluid (HTF; ART-1020) medium. Samples were treated with CXCR7-specific neutralizing antibody (ab72100, 10 μg/ml) and CXCR4 specific inhibitor (AMD3100, 2 μΜ) sequentially or at the same time, and then incubated at 37 °C under 6% CO_2_ in humidified air for 3 h. Aliquots of 10 μl were added to different concentrations of CXCL12 in HTF medium. After mixing, 5 μl aliquots were dropped onto the counter plate and analysed using a computer analysis system (WLJY-9000, Weili, Beijing). If the viability of spermatozoa in the sample was <60% it was not counted. For each sample, eight different fields were randomly selected and calculated for ≥400 spermatozoa. The sperm motion parameters recorded were VCL (curvilinear velocity), VSL (strait-line velocity), VAP (average path velocity), ALH (amplitude of lateral head displacement), LIN (linearity), and BCF (beat-cross frequency). The experiment was performed with 32 independent samples.

### Acrosome reaction

The effect of CXCL12 on spermatozoa acrosome reaction was measured by FITC-PSA (Lectin Pisum sativum(pea) as previously described ^56^. Capacitated sperm samples (25-30 × 10^6^/ml) were incubated with CXCL12 (100ng/ml) for 3 h, and then aliquots of 10 μl were fixed with 98% methanol for 10 min, and then rinsed with PBS for three times. After incubation at room temperature for 1.5 h with FITC-PSA (2 mg/ml; Invitrogen), they were washed with PBS for three times. Images of acrosome-reacted spermatozoa were captured using an Olympus fluorescence microscope at a magnification of 400 ×. Calcium ionophore A23187 (10 μM) was used as a positive control and dimethyl sulfoxide (DMSO) solvent as a negative control. The acrosome reaction rates, defined as loss of the acrosome above the equatorial segment of the sperm head, were counted for 200 spermatozoa. The experiment was performed with 6 independent samples.

### Microcapillary bioassay

The examination of human sperm penetration into viscous media (1% methylcellulose) was performed as previously described ^44^. The ends of 7.5-cm flattened glass capillary tubes (1.0-mm inner depth; Elite Medical Co., Ltd., Nanjing, China) were sealed with rubber cement. The capillaries were filled with 1% w/v methyl cellulose (V900506, Sigma) solution dissolved in HTF medium using a microinjector. Marks were placed at 1 and 2 cm on the microcapillary tube. Aliquots of 10 μl capacitated sperm samples were injected into the tubes at one end and incubated at 37 °C in a 6% incubator for 1 h. The numbers of spermatozoa reaching 1 and 2 cm were counted under a microscope. Attention was paid to avoid any bubbles during the above operations, otherwise the sperm motility would have been affected. The experiment was performed with 24 independent samples.

### Trans-well method for the study of sperm chemotaxis

The chemotaxis of CXCL12 was detected by trans-well plates (8 mm pore size, 6.5 mm diameter; Corning, NY, USA) according to the previous study ^16^. In the lower chamber of each trans-well plate, 200 μl HTF medium was added with different concentration of CXCL12 and 50 μl aliquots of capacitated sperm suspension from different treatment groups were added to the upper chamber. After incubating for 3 h at 37 °C under 6% CO_2_ in humidified air, the upper chamber was taken out. The medium in the lower chamber was mixed by pipetting and 5 μl aliquots were placed onto a cell counter (Sefi Medical Instruments-Makler) to calculate the total number of spermatozoa. The experiment was performed with 21 independent samples.

### Detection of calcium concentrations in spermatozoa

Changes in Ca^2+^ concentration in sperm cytoplasm were detected using 5 μM calcium indicator Fluo4-AM (Invitrogen) containing 0.1% Pluronic-acid F-127 and incubated at 37 °C under 6% CO_2_ in humidified air for 30 min in the dark ^57^. Redundant Fluo4-AM was removed by washing with HTF medium. Sperm suspensions (200 μl) were mixed with different concentrations of CXCL-12 in cell culture dishes and Ca^2+^ changes in motile spermatozoa were monitored using confocal laser scanning microscopy (Nikon, A1R). We used a sample frequency of 1 Hz, recorded for 3-4 min after stimulus. Changes in calcium fluorescence levels in sperm heads and midpieces were expressed as F/F_0_ ratios, where F represented the average fluorescence among treated samples and F_0_ donated the average fluorescence of the control group. Video recordings were analysed using NIH ImageJ software (https://imagej.nih.gov/ij/). The experiment was performed with 22 independent samples.

### Statistical analysis

All experiments were repeated at least three times. Results are expressed as the mean ± standard error of the mean (SEM). Differences between experimental and control groups were analysed using one-way analysis of variance (ANOVA) followed by Dunnett’s *post hoc t*-test, when appropriate. A *P* value of less than 0.05 was considered statistically significant. All the above operations were run in SPSS.

## Acknowledgements

We thank professor Jianyuan Sun (Laboratory in the Institute of Biophysics, Chinese academy of sciences, Beijing, China) for his help in the complement of the experiments and professor Zhanchun Tu (Beijing Normal University, Beijing, China) for his suggestion on the physical model.

